# Predictive learning enables compositional representations

**DOI:** 10.1101/2025.09.26.678731

**Authors:** Gauthier Boeshertz, Claudia Clopath

**Affiliations:** Department of Bioengineering, Imperial College London

## Abstract

The brain builds predictive models to plan future actions. These models generalize remarkably well to new environments, but it is unclear how neural circuits acquire this flexibility. Compositional representations, which have been observed in the brain, could explain this adaptability. They enable a process called compositional generalization, where independent modules performing different computations can be selected to perform novel composite tasks. In this work, we show that compositional representations emerge in recurrent neural networks (RNNs) trained solely to predict future sensory inputs. We trained an RNN to predict frames in a visual environment defined by independent latent factors and their corresponding dynamics. We found that the network learned to solve this task by developing a compositional model. Specifically, it had disentangled representations of the latent factors, and formed distinct, modular clusters, each implementing a single dynamic. The network autonomously selected which cluster to use according to the sensory inputs, without task labels, using competitive dynamics between clusters. This modular and disentangled architecture enabled the network to perform compositional generalization, accurately predicting outcomes in novel contexts composed of unseen combinations of dynamics. Our findings explain how an unsupervised mechanism can learn the modular causal structure of an environment in a compositional code.

**Significance:** The brain can function in environments it has never been in before, an ability called generalization. In our study, we show that when a neural network model is trained to predict future observations, it will develop a modular structure that enables the brain to do one type of generalization, called compositional generalization, where known dynamics can be recomposed in new ways. Indeed, the network develops clusters that each perform a computation representing a dynamic. Importantly, it can recombine these clusters according to the input it is receiving, with no need for other clues. We also show that the network has distinct clusters of neurons coding for each latent of the environment, consistent with what has been observed experimentally.

## 1 Introduction

To anticipate the consequences of actions or events and plan accordingly, the brain is widely believed to employ world models [13, 46, 52]. This principle is fundamental to motor control, where the cerebellum is hypothesised to predict sensory outcomes to refine movements [17, 42], and to sensory processing, where predictive coding theories posit that higher-level cortical areas send predictions to lower-level areas to be compared against incoming sensory data [5, 11, 33, 49]. These models generalise remarkably well to new environments, but it is not clear how. Here, we show that predictive learning naturally leads to a type of generalisation called compositional generalisation [20, 52].

Compositional generalisation is the ability to solve new tasks by selectively using and combining learned representations encoding elementary parts of states and rules of an environment. It is thought that the brain operates on this principle [15, 35, 40, 43, 44, 45]. For example, prefrontal networks integrate task rules with sensory inputs [35], motor systems stitch primitives into sequences [45], and hippocampo-cortical loops recombine relational structure to support inference and generalisation [40]. To explain how compositionality could arise in the prefrontal cortex, recent work has demonstrated that RNNs trained with multi-task learning can develop compositional solutions, spontaneously organising into modular subnetworks that handle distinct, elementary computations [8, 48, 53]. This modular architecture facilitates efficient generalisation, as novel tasks can be addressed by reassembling these learned components. While this paradigm is a powerful model for goal-directed learning, it is less clear how it applies to processing in sensory systems, which don’t have access to many tasks and labels.

Recent work has shown that an RNN trained with a predictive objective in an environment governed by a single latent variable learns to represent that latent in its hidden state activity [34]. However, the type of representations that would emerge in a rich environment, governed by multiple factors and dynamics, is not clear. We hypothesised that for an RNN to successfully predict future observations in such a composite world, it must implicitly learn the underlying causal structure. Therefore, training with a purely predictive objective should drive the emergence of a modular and compositional internal representation, where distinct latent factors and their dynamics are segregated in the network’s state space.

To test this hypothesis, we adapted a task used in [45] for our purposes of sequential predictive learning. [45] trained monkeys to draw a collection of shapes with different scales on a screen. We implemented this task with a dataset in which coloured shapes appear at multiple screen positions and at any temporal position in a sequence. To assess compositional generalisation, we left some compositions of dynamics that change the latents out of the training data, and tested the trained network on them.

Compositionality is closely related to a concept called disentanglement, also called neuronal selectivity in neuroscience. In machine learning literature, it has been widely studied in autoencoder models [1, 4, 14]. A model is said to have disentangled representations of the latents if, in some layer, usually a bottleneck, the latents are encoded in distinct neurons. Unfortunately, decoders receiving perfectly disentangled inputs from the bottleneck do not automatically generalise to new compositions of the latents [23, 28, 29]. Certain architectural constraints or data augmentations can help in this regard [23]. In this work, we focus on the composition of dynamics that act on latents, and not on the composition of latents themselves. This distinction is important, as it allows us to constrain where the modularity should come from. In image models, a standard decoder with deconvolutional layers needs every stage to be modular [23], whereas we only require the RNN, which computes the dynamics, to be modular. Thus, if the RNN has modular representations of the dynamics, we can expect the entire model to generalise compositionally on the dynamics. Furthermore, with dynamics encoding information about only one latent, the modular network should have disentangled representations of these latents too.

Moreover, although modularity and compositionality are closely related, they characterise distinct properties of a computational system. Modularity concerns functional organisation, different components specialise in separable computations and can be selectively engaged, reducing interference and enabling reuse across tasks[8, 27]. Modularity alone, however, does not guarantee compositionality, because a system could find the right modules for every training primitive without being able to reuse these modules in novel contexts. Compositionality instead occurs when components learned individually are recruited according to a recombination rule to solve combinations not encountered during training [20, 39]. Recent work places compositional mechanisms on a continuum defined by the expressivity of their building blocks and the complexity of the rules governing their recombination [37]. In our model, the latent-specific clusters provide evidence for modularity because each preferentially implements a single dynamic. Furthermore, performance on held-out contexts provides evidence for compositionality because dynamics encountered individually during training are selected together in novel combinations.

We trained an RNN with predictive learning in this environment. We found that the network learned to disentangle the latents, and that a modular structure comparable to that achieved through multi-task learning, whose organisation did not require explicit task or context labels. Both of these properties enabled the network to generalise compositionally.

## 2 Results

To investigate how predictive learning enables (Figure 1 A) compositional representations, we constructed a synthetic dataset of videos (Figure 1 B), in which each frame is uniquely determined by three discrete independent latent variables, which encode the object’s shape (*z_shape_*), colour (*z_colour_*), and position (*z_position_*) as their index in their respective list of possible values. We generated sequences from these images by applying dynamics that independently change one latent. Specifically, each sequence is associated with a context, defined as a tuple of offsets (Δ*z*_shape_, Δ*z*_colour_, Δ*z*_position_) which specifies how the corresponding latents evolve across time steps. Their composition forms the context of the sequence. For example, the context (3, 1, 2) increments the shape index by 3, the colour by 1, and the position by 2 (Figure 1 C). We used 6 dynamics for the shapes, and 3 for both colour and position. This resulted in 54 context, which we split into 43 training contexts and 11 generalisation contexts. The generalisation split has compositions not seen during training, but their individual components were. If the network performs as well on the generalisation as it does on the training set, we say it generalises compositionally.

**Figure 1:**
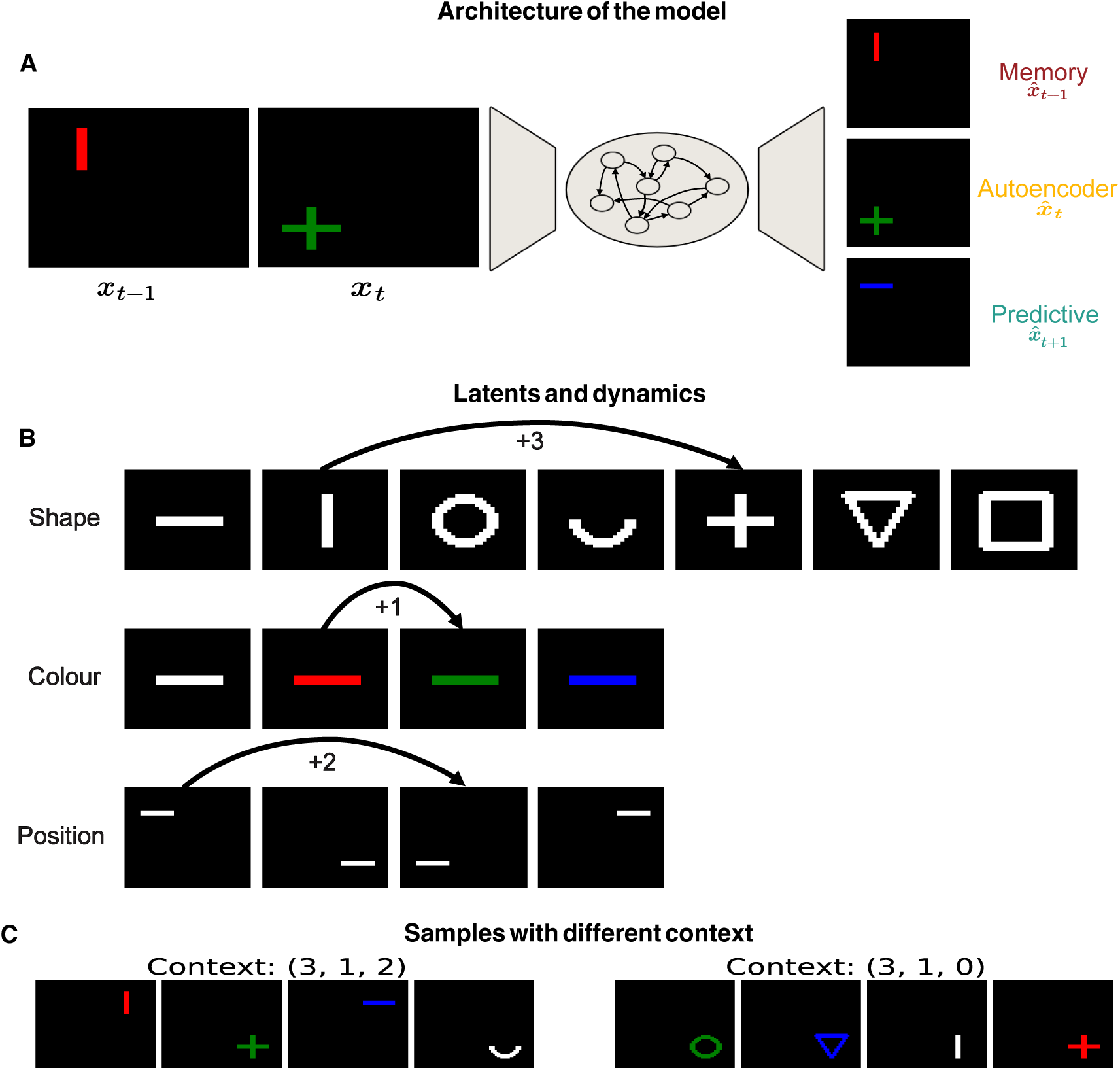
Training a network to predict the next observation. **A** The predictive model in green outputs what will be the next observation, the autoencoder outputs the last image it received and the memory model outputs the previous image. **B** We defined a number of shapes, colours, and positions. On top, we defined dynamics for each latent, that each increment by some value the index of its associated latent in its list of possible values. These lists are circular, so the addition goes around the end to the beginning. The dynamics are fixed per sequence. We call the composition of these dynamics a context. **C** Example of sequences with different contexts.

The whole network is made of a convolutional encoder that processes each image individually; the encodings are then sequentially given to the RNN, which will predict the dynamics. Its activity is fed to a deconvolutional decoder, which reconstructs the image. Importantly, it does not receive any information on the current context, but the network can infer it by integrating two consecutive inputs. The network is trained to output images directly, we do not impose any constraints beyond regularisation on weights and activations. This means that the network is free to represent the inputs in any manner. However, certain representations could make it easier for the network to generalise compositionally. In particular, having disentangled representations with respect to the latents and having clustered networks that perform the dynamics may help the network capture the underlying structure of the data. In the following, we will show how the predictive model exhibits both properties. We will compare it to an autoencoder, a network with the same structure but trained to output the last observation it received.

### 2.1 The predictive model disentangles the latents

In our modular environment, where latents change independently from each other, faithfully representing these dynamics implies that the latents will be disentangled. Therefore, before testing whether the network exhibits modular representations of the dynamics or not, we measured the disentanglement of the latents. The other way around, however, is not necessarily true; disentangled representations of one latent do not imply that the dynamics of this latent will be disentangled. For example, the representations of every dynamic for a latent could be entangled with one another; having this for every latent would make them disentangled, but would not result in modular dynamics.

To assess the separability of the latents in the network activity, we developed a metric inspired by [2, 16] (Figure 2 A). For a given target latent, we trained a regularised classifier to predict that latent from the RNN activity, and evaluated how it generalises across changes in the distribution of the non-target latents. Specifically

**Figure 2:**
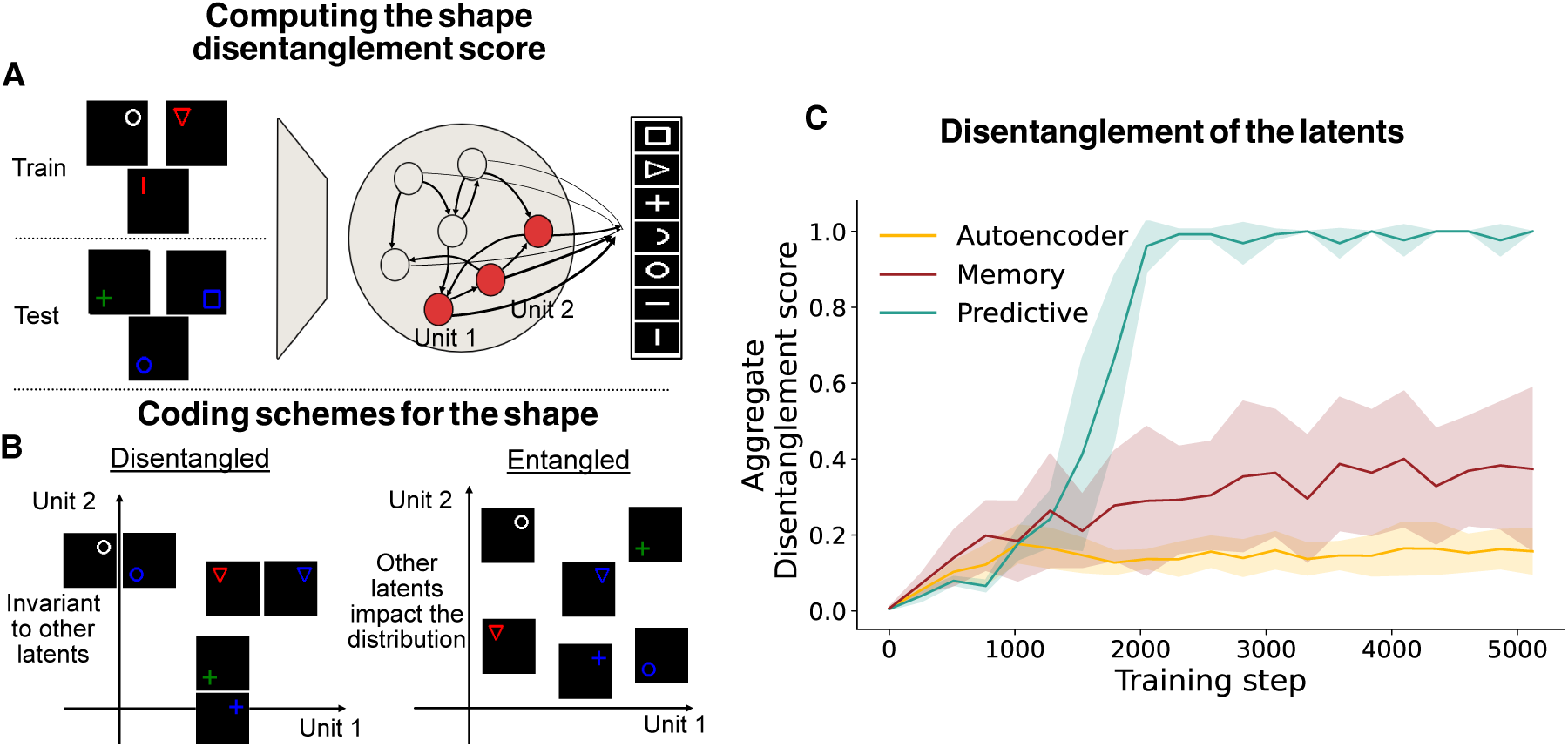
The predictive model perfectly disentangles the colour, shape, and location latents. **A** Computing the disentanglement of the shape latent. We split the support of the two colour and position latents in half, then encode the images of both splits. The training set has white and red shapes in the top quadrants. The test set has green and blue shapes in the bottom quadrants. We then train a classifier, and its performance on the test set is the disentanglement score. **B** Modes of distribution of the neurons the decoder uses. If the model has disentangled representations of the latent, the distribution is invariant to the colour and position latents, and will not change across train/test splits. Therefore, the disentangled models will have a high decoding accuracy. Entangled models will have different distributions across splits, so the decoder will perform poorly on the test set. **C** Disentanglement performance of the autoencoder, memory, and predictive models. The predictive model disentangles the three latents, while the representations of the other two are entangled.

### 2.2 The predictive model has clustered dynamics representations

Having distinct populations that each compute one dynamic could help the network be compositional, as it enables it to choose which population to use according to the information it is integrating. In contrast, there are two other ways to solve the training task that could work but would not lead to compositional generalisation. The first is having a cluster for each context encountered during training, the second is having a single, large population in which all dynamics are intermixed. In the first case, the network effectively memorises context-specific solutions rather than learning reusable components, which limits generalisation to unseen contexts. In the second, the lack of specialisation forces all computations to be shared across latents and contexts, making it difficult to isolate and recombine specific dynamics. While both approaches could achieve low training error, neither would produce the compositional structure that would generalise to unseen contexts.

To test the compositional properties of the network and assess which of the solutions explained above the network adopted, we measured the degree of clustering of the representations using a method from [53]. Specifically, we first computed the normalised variance of each neuron across contexts, which serves as a measure of context selectivity. These selectivity profiles were then partitioned into clusters, with the number of clusters chosen to best explain the data. We computed the number of clusters for the predictive model (Figure 3 A) and for the autoencoder (Figure 3 C) and sorted them according to the latent they affected. We found that the predictive model had clear clusters that each perform one dynamic. The autoencoder and the memory models did not have clusters because they did not need to compute any dynamics. To understand the causal role of the clusters, we computed (Figure 3 B) the difference in loss when lesioning one cluster of neurons from the original loss. The differences are similar to the selectivity measure, confirming that a cluster is only active when its selected context is active. Furthermore, removing a cluster from the network impacts only the aspect of the object related to the dynamic computed by the cluster (Figure 3 E). For example, cluster 4 changes the shape of the object; lesioning it affects only the shape of the object in the output. Indeed, the position and colour are both correct, but the shape looks like the mean of all shapes.

**Figure 3:**
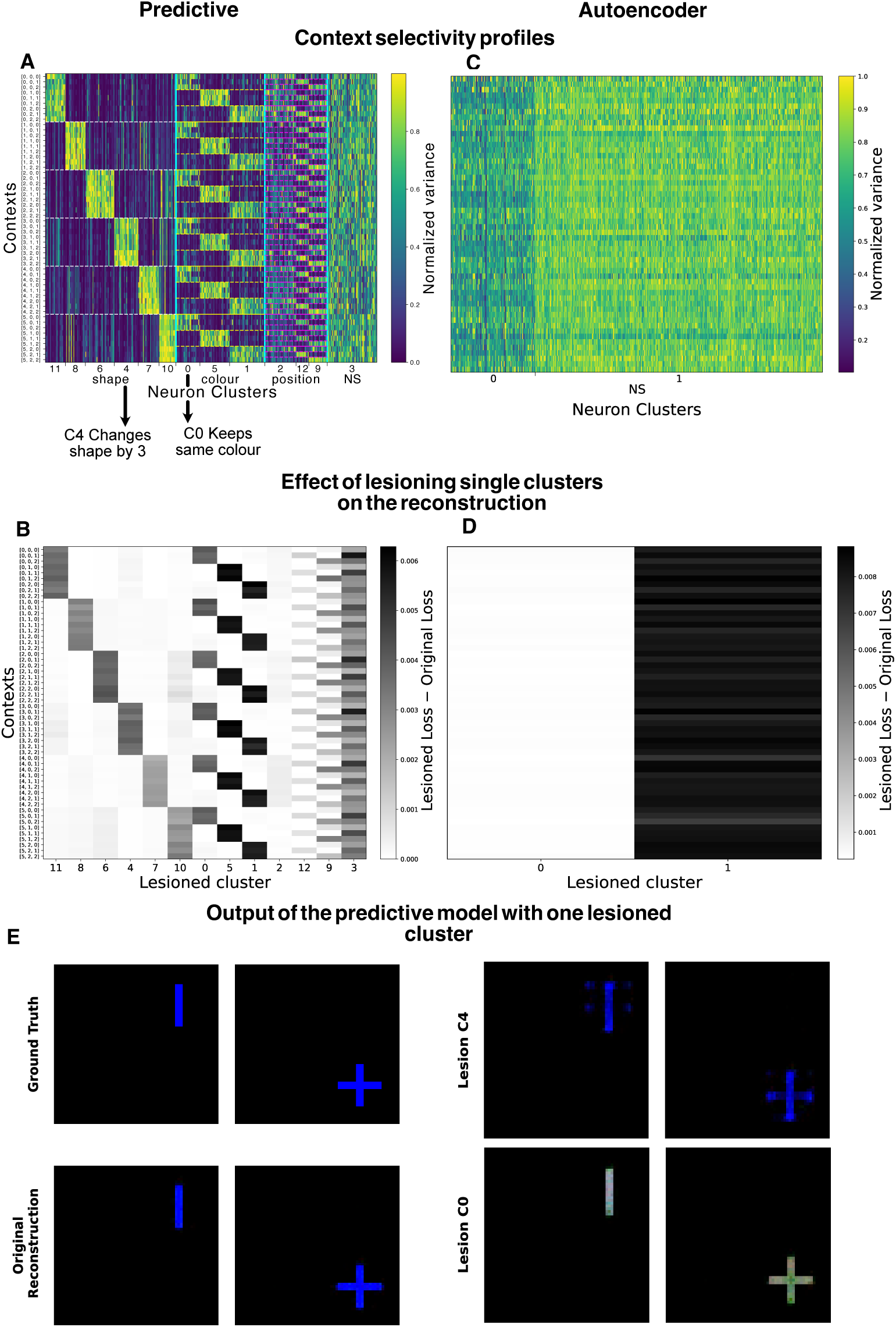
Clustered dynamics in the predictive model. **A** Sorted context selectivity of the predictive model. Each dynamic has its own cluster that represents it. For example, cluster C4 codes for the sequence that changes the shape by 3 each transition, and cluster C0 codes for the sequence where the colour doesn’t change. We marked each group of clusters by the latent it affects, and NS for cluster not selective to any latent. **B** The effect of lesioning a cluster on the predictive model’s output is largely constrained to the context the cluster is selective for, as measured by the difference in loss between the lesioned model and the original. **C** Same as **A** but for the autoencoder. There are only 2 active clusters. **D** Same as **B** for the autoencoder, where lesioning cluster 1 affects every context. **E** Outputs of the predictive model when lesioning the relevant cluster to the context. Lesioning C4 results in a degenerate shape that does not resemble any shape. For cluster C0, the effect is also isolated to the colour.

### 2.3 Mechanism of cluster selection

Having established that the predictive model develops clustered dynamics, we investigated the neural mechanisms that enable these properties. First, we found that the network employs a selective activation mechanism. When processing an input sequence, only the clusters selective to the sequence’s context show significant activity, while the others show little activity(Figure 4 A, B). The network organises itself according to the context at hand, and each cluster inhibits the other clusters relevant to the same latent. To understand how the clusters are selected, we computed the cosine similarity between the input and recurrent current within a cluster for every cluster computing the dynamics of the first latent (Figure 4 B). The cosine similarity measures the angle between two vectors by taking the dot product of the normalised vectors. We found a spike in similarity that is always the biggest for the cluster selective to the context. In other words, the chosen cluster is the one where the input coming from the encoder and the recurrent input have most similar directions. Moreover, this selection is also aided by a competitive process between clusters associated with the same latent variable. We observed that the RNN’s recurrent weights between competing intra-latent clusters are strongly negative, while recurrent weights within each cluster are positive (Figure 4 D). This connectivity pattern is consistent with winner-take-all dynamics, where the contextually appropriate cluster becomes active, and the others are shut down. Moreover, connections between clusters coding for the same latent are stronger than connections between clusters coding for different latents.

**Figure 4:**
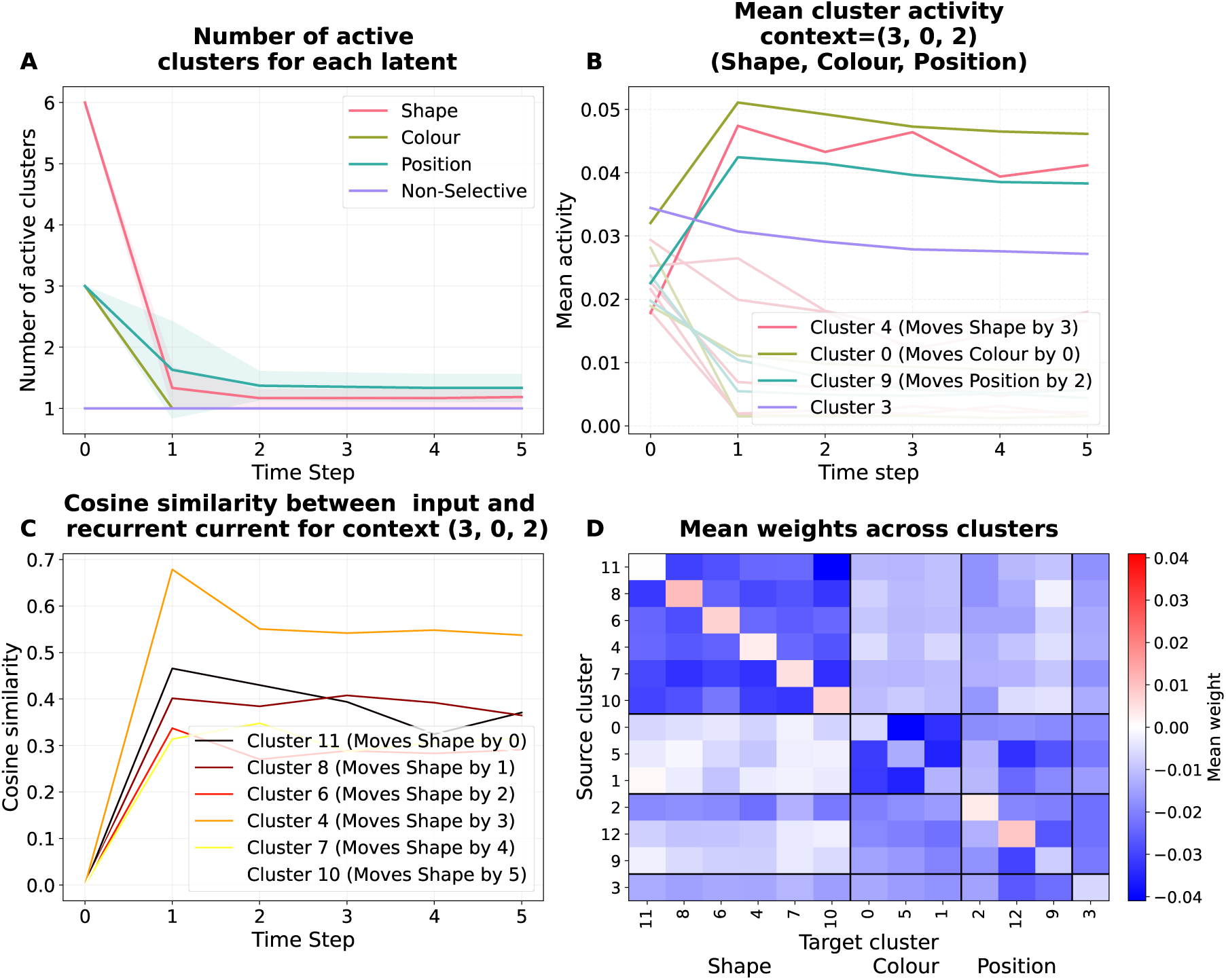
Cluster selection in the predictive model. **A** Average number of active clusters coding for each latent in every context. On average 1 cluster is active for each latent. We defined active as clusters whose mean activity is more than half of the maximum mean activity of each clusters. **B** Example of mean cluster activity for a sequence belonging to context (3,0,2). The clusters selective to the context, for example C0 and C4, exhibit the highest mean activity out of all the clusters coding for their latents. **C** Cosine similarity between input current and recurrent cluster by clusters, for sequences in context (3,0,2). Cluster C4 has the highest similarity, and is the one that is most selective to the context. **D** Mean weight across clusters and intra-cluster. The weights between clusters coding for dynamics of one latent are more negative than those of clusters across latents.

### 2.4 The predictive model generalises compositionally

Systematic compositionality is defined as the capacity to assemble a finite set of known elements into any novel combination [7, 10, 20]. In our work, these elements are the dynamics of the latents, and we say that a model generalises compositionally if it performs as well on compositions of components not seen during learning as it does on the compositions it was trained on. We train the network on 80% of the possible compositions of the dynamics during training and save the rest for testing this compositional generalisation (Figure 5 A). We computed the training and generalisation loss (Figure 5 B), and found that the two losses go down in unison, where the generated images match the ground truth. We concluded that the model composes its clusters, using only sensory inputs, to accurately predict the next observations, and that it can do so in contexts not seen during training. We stress that it does not require any task labels, and can select which cluster to use autonomously, as opposed to common multi-task learning procedures, which give information about the current task to the network [8, 36, 53]. We also found that this ability emerges at around the same time during training as when the model perfectly disentangles the latents, as shown by the product of the score of disentangling the three latents, and when the number of clusters increases (Figure 5 B). This supports the fact that the two properties described above help the network generalise compositionally.

**Figure 5:**
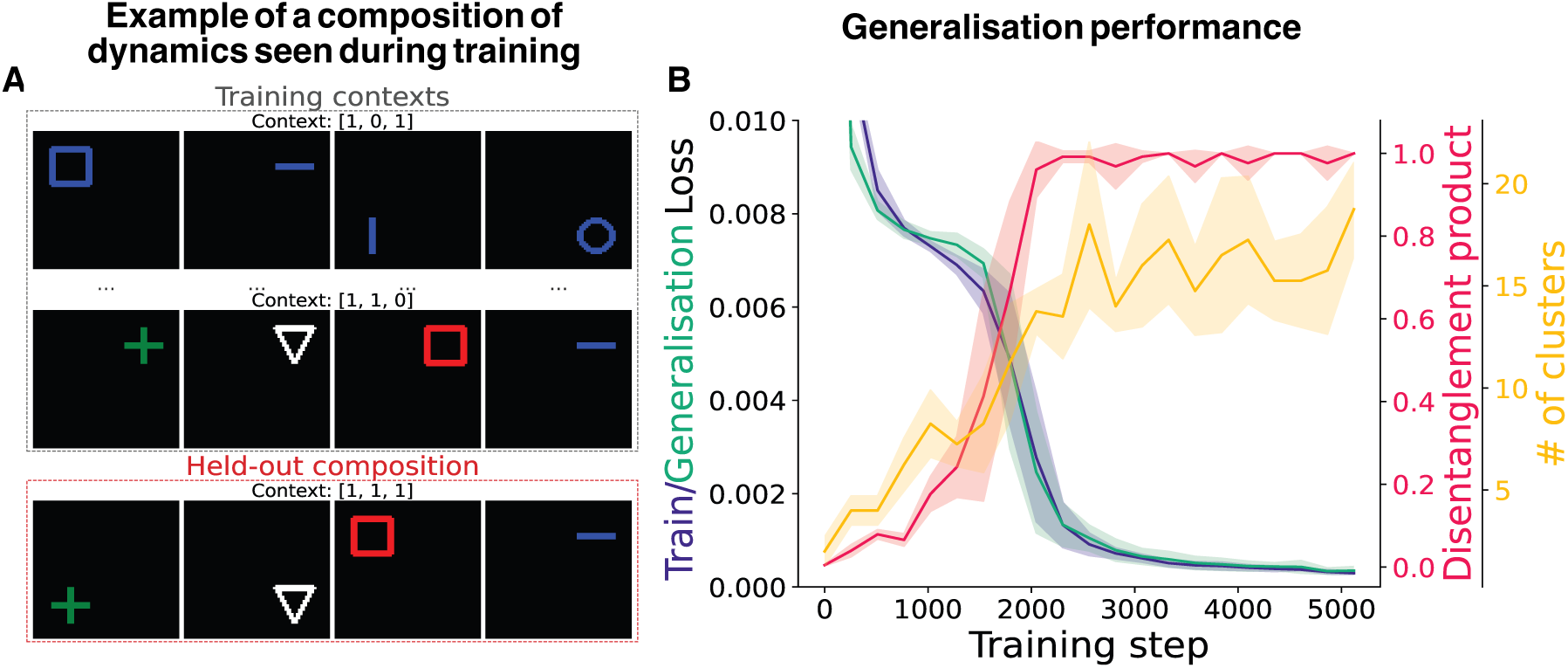
The predictive model generalises compositionally. **A** We held out some of the composition of the dynamics during training time to assess compositional generalisation. **B** The model accurately reconstructs sequences with contexts not seen during training, at a time that coincides with the disentanglement and the number of clusters reaching their limit.

## 3 Discussion

We have demonstrated that an RNN trained with predictive learning in a rich environment generalises compositionally by disentangling the generative factors and composing the dynamics. The model trained to predict the next observation separates all latents perfectly, each being encoded in distinct groups of neurons, whereas an autoencoder has them entangled. The predictive model also has multiple clusters of neurons performing one dynamic on one latent, which can be composed according to the dynamics the inputs follow.

Our work builds upon previous studies showing that RNNs trained with multi-task learning can develop modular, compositional solutions [8, 36, 53]. However, a key distinction of our model is that this structure emerges autonomously from a single objective, without the need for explicit task labels or context signals. This is particularly relevant for sensory systems, which must learn the structure of the world largely through unsupervised observation, rather than through a curriculum of distinct, labeled tasks [12, 26]. Our results reinforce the idea that predicting future sensory input can lead to informative representations, and could be used in the brain [6, 9, 12, 21, 34, 51]. Notably, predictive learning is also the principal training objective in large language models [32, 47], typically instantiated as next-token prediction, showing the generality of this learning paradigm.

The emergence of disentangled representations of the latent factors is a cornerstone of this process, as it allows for simple dynamics that can easily modify one latent only. Such invariance is one of the necessary properties of one type of compositional representation, called symbolic representations, which have been observed in the monkey brain [45]. In the direction of separability and selectivity, we predict that neural networks trained with predictive learning will develop clusters of very selective neurons that react to distinct factors of the inputs, similar to what is learned with multi-task learning in feedforward neural networks [16]. On the other hand, our simple autoencoder developed mixed neurons. More complex autoencoders, models with probabilistic latent spaces like variational autoencoders (VAEs), have been shown to learn disentangled representations [14], and have responses similar to those of biological circuits [14]. However, this success often depends on tuning the objective function to explicitly penalise disentanglement in the latent space, thereby encouraging factorisation. Other methods building on top of VAEs also use temporal information by defining transitions between images [19, 22, 24, 38], but they have not analysed simple RNNs with only a predictive objective. Our findings present a complementary perspective where temporal prediction is sufficient to drive the emergence of a disentangled code. The failure of our standard autoencoder to disentangle these factors highlights that merely reconstructing the present is not a strong enough constraint; the pressure to anticipate the future is what drives the network to learn the underlying causal structure of the environment. In addition to informative representations, learning the dynamics also allows planning and simulation of the environment.

Mechanistically, our model selects its clusters through competition between clusters impacting the same latent. These clusters compete via inhibitory connections, creating winner-takes-all dynamics where the active cluster is determined by the ongoing sequence context. This allows the network to flexibly select and apply the correct dynamic transformation while leaving other, irrelevant latent representations untouched. This functional segregation is further supported by our connectivity analysis, which reveals that clusters are preferentially connected to the neurons encoding the specific latent they manipulate. This suggests the formation of distinct processing pathways, characteristic of modular computation in artificial and biological neural networks.

Our work opens several research directions. Our synthetic environment is deterministic and relatively simple. Future studies should investigate whether these principles scale to more complex, naturalistic environments with stochastic dynamics and hierarchical structure. It would also be interesting to study how changing the environment during learning affects the network. For example, one could add new factors, such as a scale to the objects, and test whether the network keeps its representations and adds new ones for the new latents. Also, when changing the space of the latents, would a new color be encoded in the same neurons as the ones encoding the other colors? It would also be valuable to explore how these learned representations could be leveraged by a downstream decision-making network, linking predictive learning directly to goal-directed behavior.

Altogether, our results demonstrate that the simple objective of predicting future states is a powerful engine for building compositional world models. This process autonomously discovers the underlying generative factors of an environment, segregates them into disentangled representations, and learns their dynamics in a modular fashion. These findings provide a mechanistic account of how the brain might develop world models that support flexible generalisation.

## 4 Methods

### 4.1 Network Architecture

As stated in the results section, the whole network consists of a convolutional encoder, an RNN, and a deconvolutional decoder. The encoder has 3 convolutional layers with a channel size 16 with ReLU activations, followed by a linear layer with a size of 512. It then projects into the RNN, which also has size 512. The decoder has the same structure as the encoder, but mirrored. The RNN update rule is:

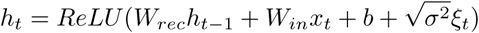

where *h_t_*is the activity of the RNN at time t, *W_rec_* are the recurrent weights, *W_in_* the inputs weights, *x_t_* the inputs, *b* the biases, *σ*^2^ is a variance scaling applied to the independent gaussian noise *ξ_t_* with mean zero and unit variance sampled every time step.

### 4.2 Training the model

We trained the predictive model to minimize the squared error of the decoder output, *ô*_*t*_, and the observation of the next-time step 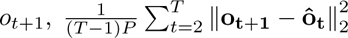, where T is the length of the sequence, and P is the number of pixels in one image. The evaluation starts from the second frame (t=2) because the model requires at least two frames to infer the sequence’s dynamics. We add L2 regularization on the weights and activity of the RNN, with weights *λ_weights_* and *λ_act_*. We didn’t find stereotypical effect of increasing or lowering these parameters, similar to [53]. We trained the whole network for 5000 optimization steps, using the AdamW [25] optimizer with a learning rate of 0.001, and momentum parameters *β*_1_ = 0.9*, beta*_2_ = 0.999, a batch size of 128, using the Pytorch library [30].

### 4.3 Hyperparameter search

We ran a grid search on the two regularization terms, both with values [0, 0.001, 0.01, 0.1, 1, 10], and size of the RNN with values [128, 512] for the predictive model and [8, 32, 128, 512] for the autoencoder and memory model. We used *λ_weights_* = 0.001*, λ_act_* = 0.01 and a size of 512 for the RNN for the predictive model because they gave the cleanest clusters. We also show the results with the lowest validation loss in the supplementary S4, with parameters *λ_weights_* = 0.1*, λ_act_* = 0. For the autoencoder, we found that *λ_weights_* = 1*, λ_act_* = 0, and size 512 achieved both the lowest generalisation loss and the best disentanglement product. For the memory model we found *λ_weights_* = 1*, λ_act_* = 0.01, and size 512. We ran the search with 2 seeds for each combination of hyperparameters, and then ran the selected hyperparameters with 4 seeds to make Figure 2 and Figure 5. The plots for Figure 3 and Figure 4 where made with the seed 0 of these models.

### 4.4 Dataset

The dataset is made of sequences of frames of size 64×64. Each is fully determined by three discrete independent latent variables, which encode an object’s shape (*z_shape_*), position (*z_position_*), and colour (*z_colour_*), as their index in their respective list of possible values. There are seven possible shapes: a wide rectangle, a tall rectangle, a circle, a semi-circle in a U-shape, a cross, a triangle, and a square. There are four possible colours: white, red, green, and blue. There are 4 positions, the center of the four quadrants of the frame. To create a video, we sample one value for each latent for the initial conditions and then apply independent dynamics to them, which we call the context of the sequence. They are defined by a tuple (Δ*z*_shape_, Δ*z*_position_, Δ*z*_colour_) For example, the context (1, 0, 2) increments the shape index by 1, leaves the position unchanged, and increments the colour by 2. We used 6 dynamics for the shapes, and 3 for both colour and position. This resulted in 54 dynamics, which we randomly split into 43 training contexts and 11 testing contexts, while making sure that every primitive was in at least one training context.

### 4.5 Disentanglement

For each target latent z, we trained a regularized linear classifier to predict z from the RNN population activity and quantified how well this decoder generalises across changes in the distribution of the nontarget latents. Concretely, we constructed disjoint training and testing splits by partitioning the support of all non-target latents into two halves, sampling frames from one half for training and from the other half for testing, while keeping the target latent’s support unchanged. When used as a nontarget latent, for the shape, we used the horizontal line, the vertical line, the circle and the semi-circle for training, and the cross, triangle, and square for testing. For the position, the top right and top left for training, and the bottom left and bottom right for testing. For the colour, the white and red for training, green and blue for testing. For example, when probing disentanglement of the shape, the decoder is trained on shapes appearing only with specific colour–position combinations, white/red in the top half of the frame, and evaluated on held-out combinations, green/blue in the bottom half. We then presented frames from both splits to the model, extracted the corresponding RNN activity, and fit a linear classifier with an L1 penalty of weight 10 to decode z, using scikit-learn [31].

### 4.6 Cluster analysis

We first calculated a context selectivity score for each RNN neuron, following the task variance analysis method from [53]. We computed the variance of each neuron across all contexts and discarded any with a summed variance below 0.001. The activity of the remaining neurons was then normalized by their individual maximum variance across the contexts.

Using these selectivity scores, we clustered the neurons with the K-Means algorithm, using the scikit-learn library [31]. To determine the optimal number of clusters, we performed a sweep from 2 to 25 clusters, selecting the configuration that minimized intra-cluster distance while maximizing intercluster distance. We then sorted the clusters based on the latent dynamic that elicited the strongest response.

### 4.7 Cluster selection

To compute how many clusters coding for the same latents are active, we counted the number of clusters with an average activity greater than half of the most active cluster’s average activity, among clusters representing the same latent variable:

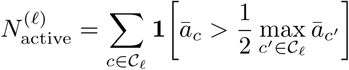

where *C_ℓ_* denotes the set of clusters coding for latent *ℓ*, and *a̅_c_* is the mean activity in cluster *c*.

Cosine similarity measures how aligned two vectors are. It is defined as the dot product of the two vectors normalised by their lengths:

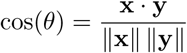

A value close to 1 indicates that the vectors point in similar directions, while a value close to 0 indicates that they are mostly unrelated. In our case, we use cosine similarity to compare the direction of the input coming from the encoder with the direction of the recurrent input.

## Acknowledgements

This work was supported by Wellcome Trust 200790/Z/16/Z, Simons Foundation 564408 and EPSRC EP/R035806/1.

---

**Figure S1:**
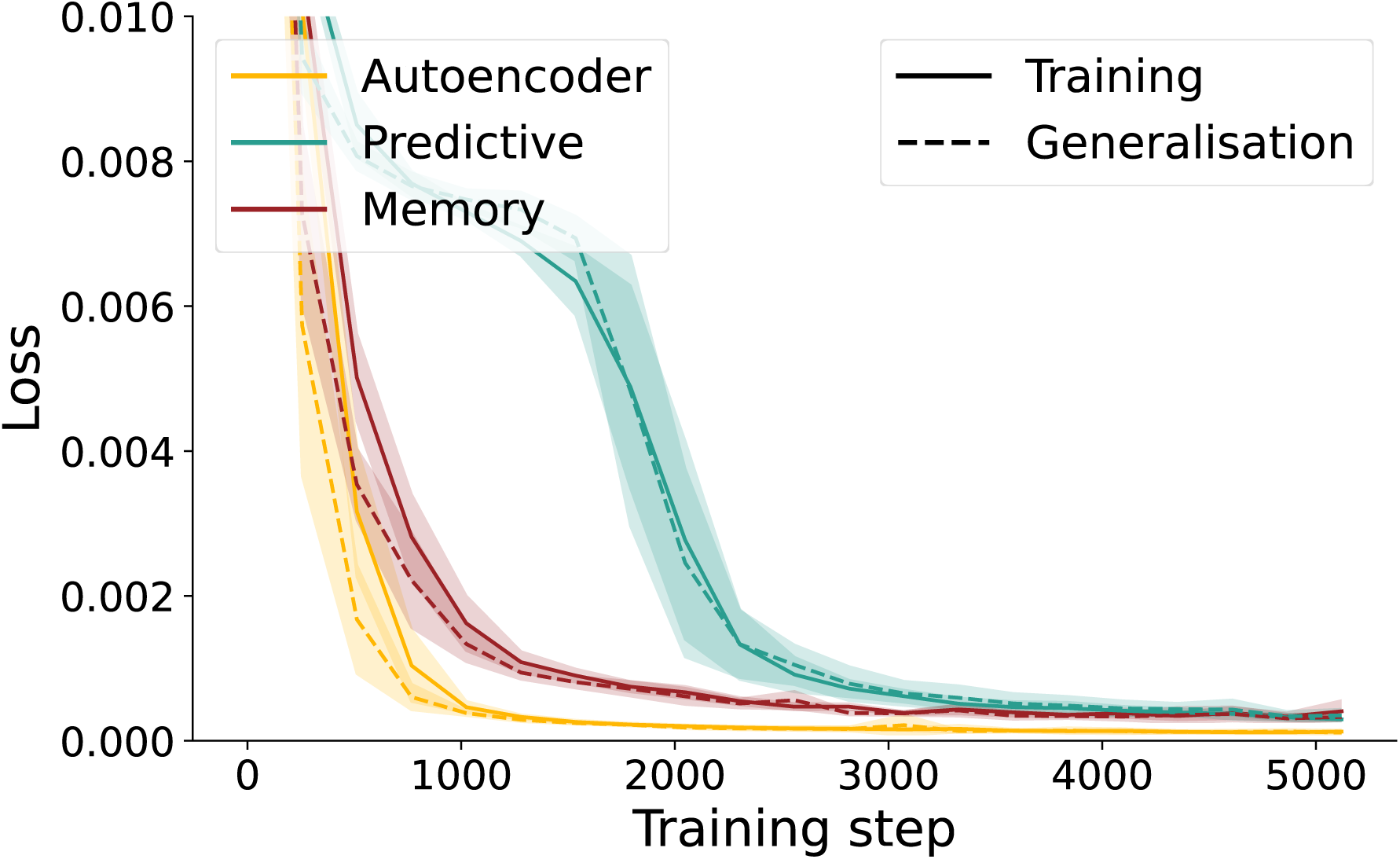
Comparing the training performance of the three models.

## Supplementary

### S1 Training performance

We computed the training and generalisation losses every 256 gradient steps during training (Figure S1). The autoencoder losses converge the fastest, the memory model soon after it, whereas the predictive model is slower. This is due to the fact that predicting the future observation is much harder than reconstructing already seen observations. Additionally, as the autoencoder and memory models do not have to take into account dynamics, training and generalisation can be seen as the same task.

### S2 Comparing the disentanglement scores across hyperparameters

To try and get the best disentanglement score for the three models, we ran hyperparameter sweeps with different sizes of bottlenecks, again following [50], with different levels of L2 regularisation on both weight and bottleneck activity (Figure S2). Most of the predictive models that generalise have disentangled latents, whereas autoencoder and memory models never disentangle the three latents and have a much broader distribution.

### S3 Comparing to an autoencoder without an RNN in the middle

Having an RNN in the middle of the encoder and decoder may affect the disentanglement. Therefore, to get another control, we computed the disentanglement score of a feedforward autoencoder with no recurrency (Figure S3). It will not use temporal information, which is fine since it only reconstructs the observation it is receiving. We ran hyperparameter sweeps with different sizes of bottlenecks, again following [50], with different levels of L2 regularisation on both weight and bottleneck activity. The RNN autoencoder disentangles the latents similarly to the CNN autoencoder.

**Figure S2:**
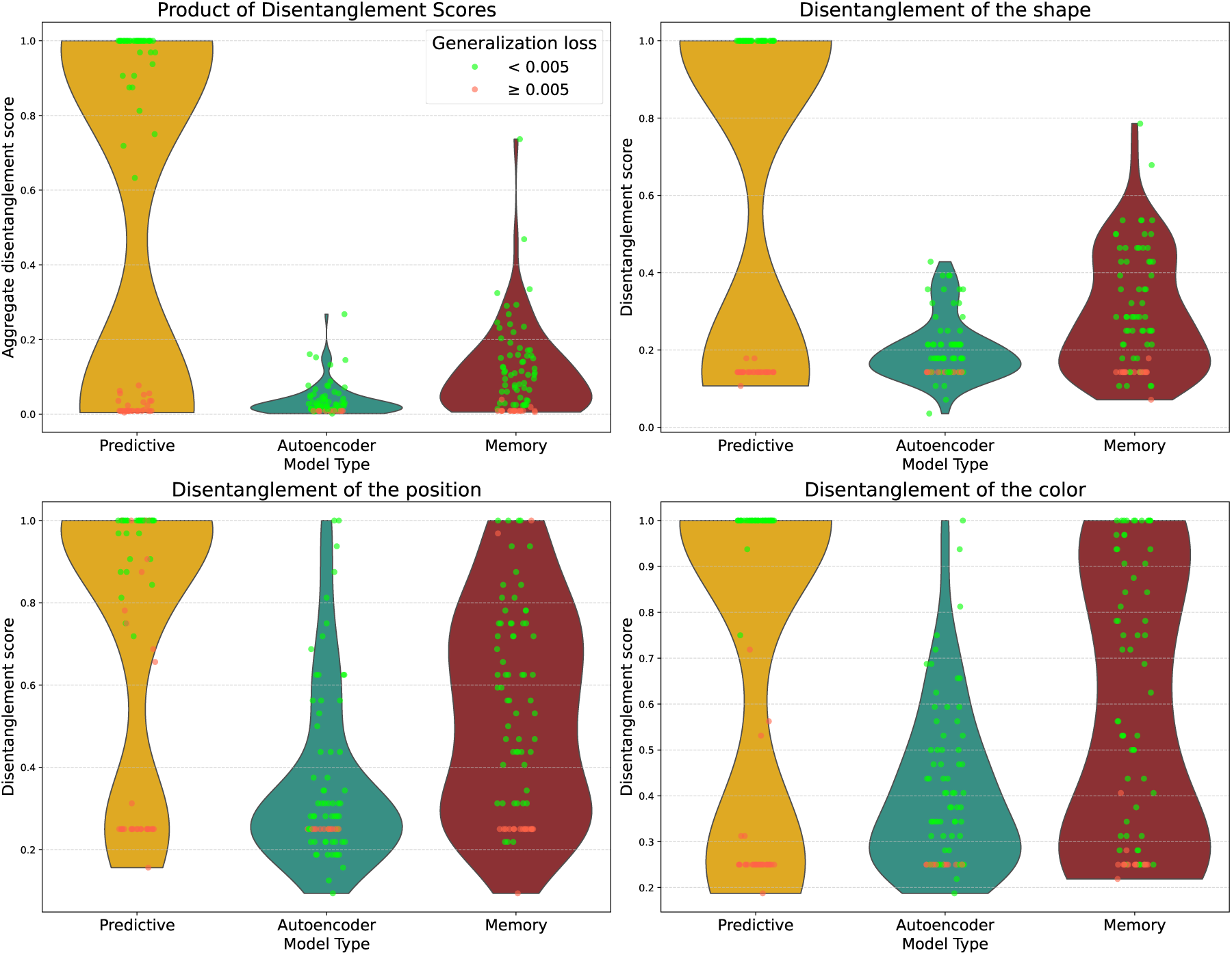
Comparing the disentanglement of the predictive, autoencoder, and memory models. Each dot is the result of one seed. The colour of the dots indicates whether the generalisation loss of the model was lower than 0.005, which we defined as a successful run. For the predictive model, a successful run almost always means that the three latents are perfectly disentangled, whereas neither the autoencoder nor the memory models perfectly disentangle all of the latents. There are fewer points for the predictive model because lower sizes of bottlenecks were never successful, so we didn’t include them in the hyperparameter search.

**Figure S3:**
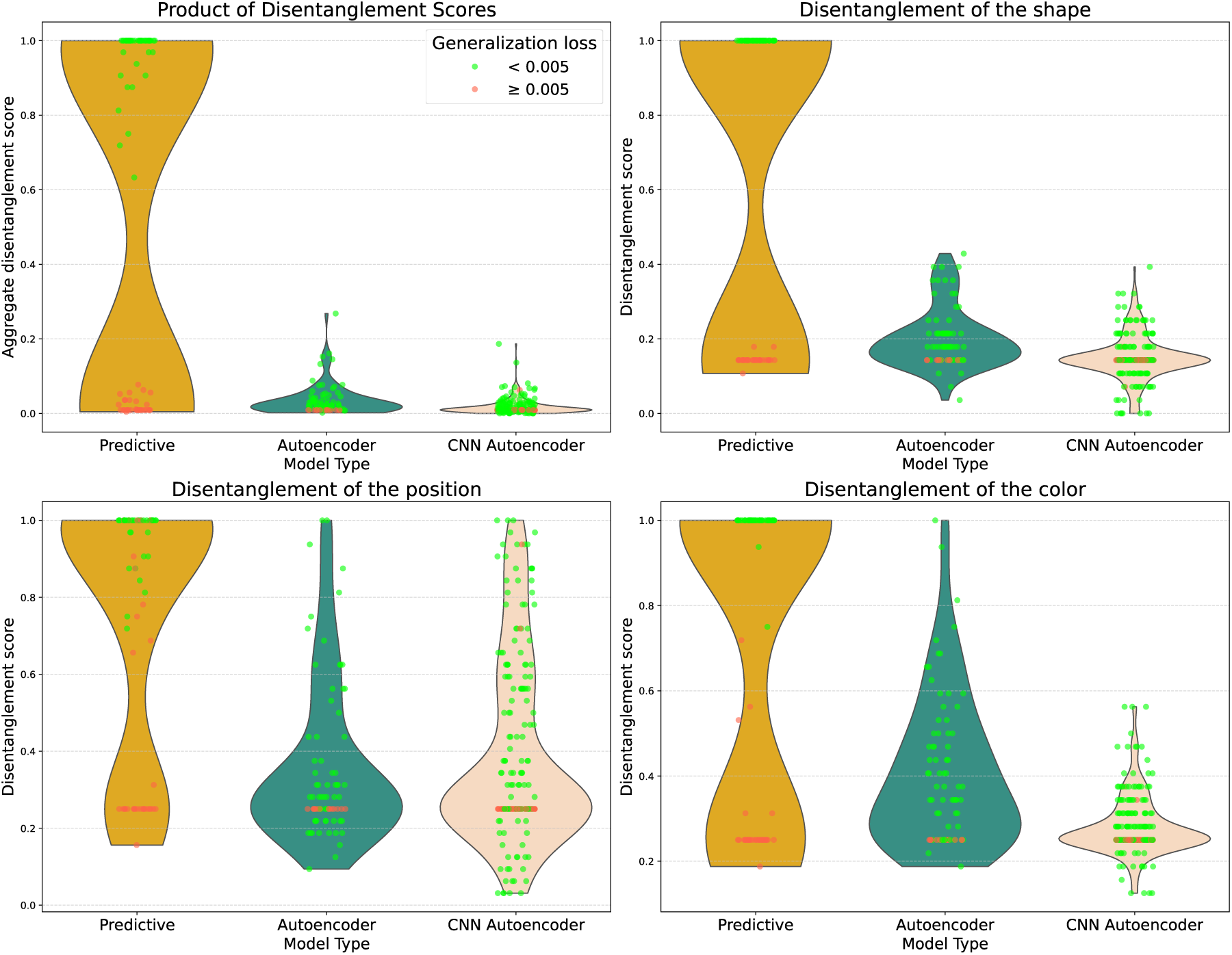
Comparing the disentanglement of the predictive model, a recurrent autoencoder, and a feedforward autoencoder. Disentanglement is usually studied in CNNs, thus we wanted to know whether using an RNN could be what makes the autoencoder not disentangle. It is not the case as the two autoencoders have similar disentanglement.

### S4 Results for the lowest generalisation loss model

**Figure S4:**
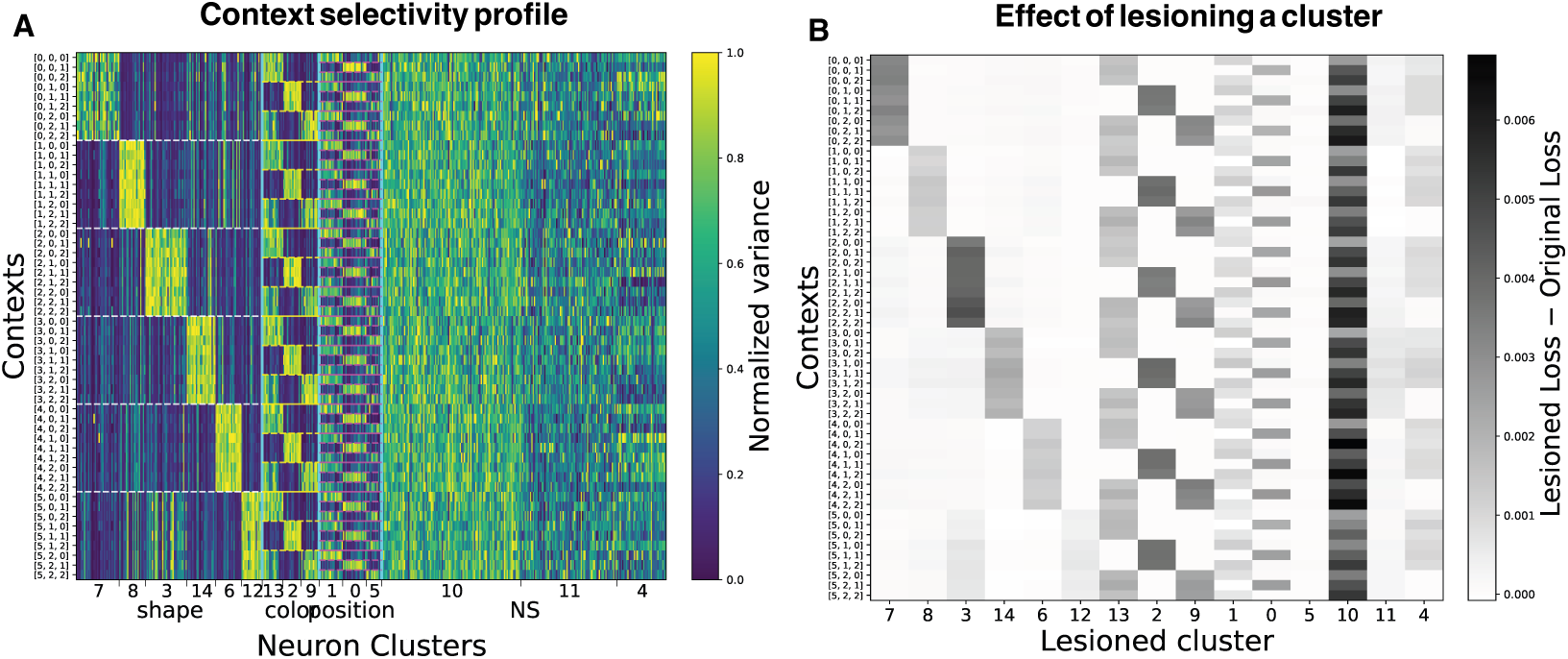
Results for the model with lowest generalisation loss. **A** Clustering analysis of the model. The model still has distinct clusters for the dynamics. **B** Effect of lesioning a cluster. Even though the model has distinct cluster for the dynamics, lesioning a cluster also has effects on contexts it is not selective to. For example, lesioning any cluster computing dynamics for the shape worsens the reconstruction for contexts associated with moving the shape by 5, even though they are not selective to it.

We chose the model with the clearest cluster for the main results, and selected the model with the lowest generalisation loss for the autoencoder, which coincided with the best disentanglement as well. We also computed the clustering and the summary statistics of the correlation between clusters and mean weights across clusters for the predictive model with the lowest generalisation loss Figure S4. We still find a clear cluster structure (Figure S4 A), but lesioning them has a greater effect in contexts they are not selective to (Figure S4 B).

### S5 Effect of using fewer tasks

The representations we analysed may result from a trade-off in the capacity of the model against the difficulty of the task. To test that idea, we also trained a model on fewer contexts by decreasing the number of dynamics per latent to 2 (Figure S5). We found that the model does not generalise compositionally, achieving 0.0003 MSE on the training contexts and 0.0018 on the testing contexts (Figure S5 A). The model does not disentangle the latents (Figure S5 B). While the model has modular representations of the dynamics acting on the shape, the rest are not as clear (Figure S5 C, D). We used the same hyperparameters as the ones used for the main text model. In conclusion, using more regularisation, or simply using fewer neurons to limit the capacity of the model may be necessary to achieve representations suitable for compositional generalisation.

### S6 Example of trajectories encoded by a cluster

The clusters each compute a sequence that describes the trajectory of one latent. We projected the activity of cluster C4, which changes the shape latent by 3 at each step, and C0, which computes the sequence where the colour does not change, using Multi-Dimensional Scaling (MDS) [41] when varying the initial value of the latent (Figure S6. The first projected activity is not the center of the other activity with the same input since the RNN has not given recurent input yet; however, right after it, the trajectory goes in a trajectory that follows the structure of the sequence we imposed. The activity of the RNN is well separated across different values, but close across different timesteps with the same latent value, which allows the decoding of the right value from the cluster.

**Figure S5:**
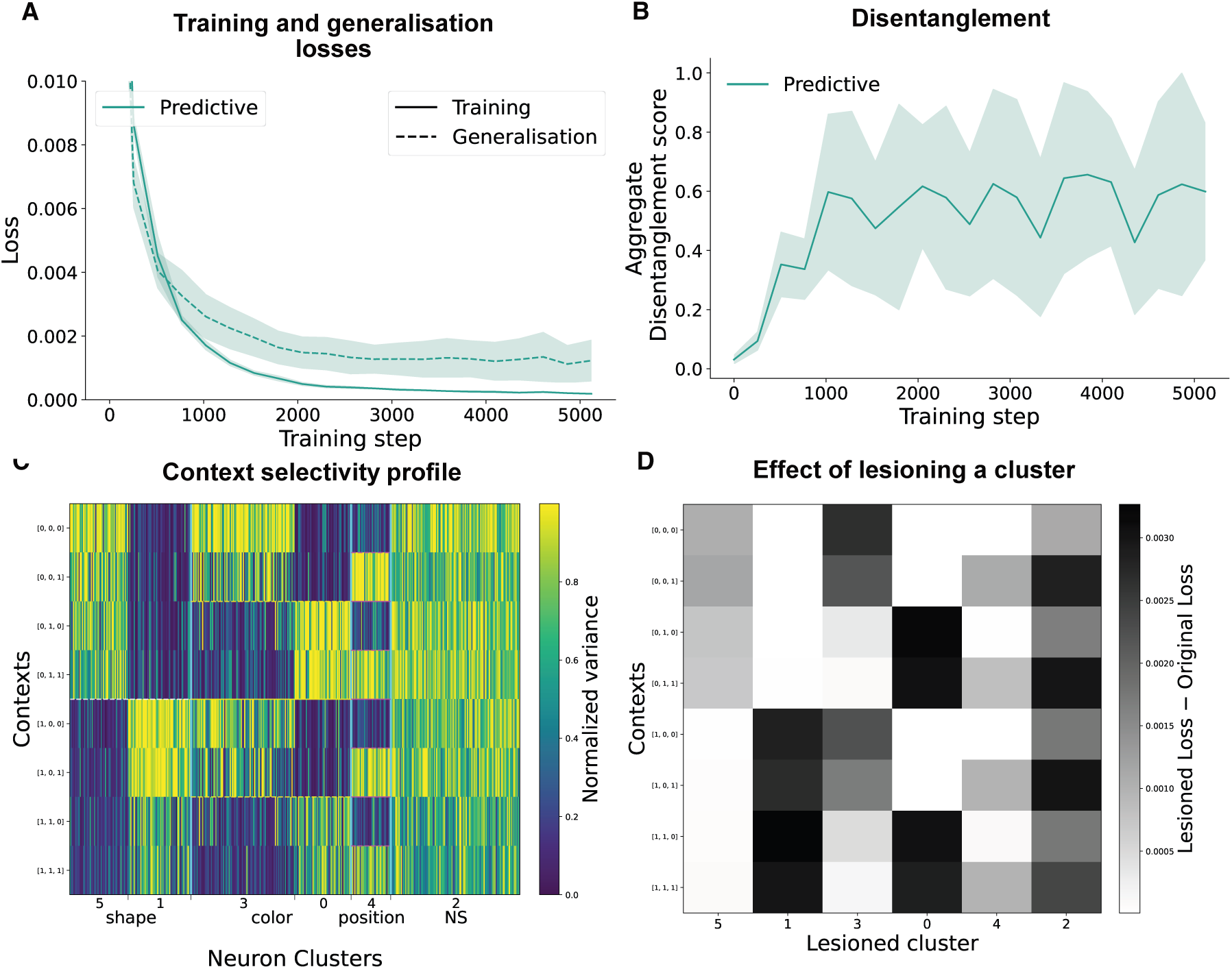
Cluster analysis of the predictive model trained on fewer contexts. **A** Training and generalisation loss. The model does not generalise to new contexts. **B** The model does not disentangle the latents. **C** Cluster selectivity. The network has modular representations of the shape dynamics, but the other dynamics are not represented. **D** Effect of lesioning a cluster. Lesioning a cluster affects the reconstruction performance across every context.

**Figure S6:**
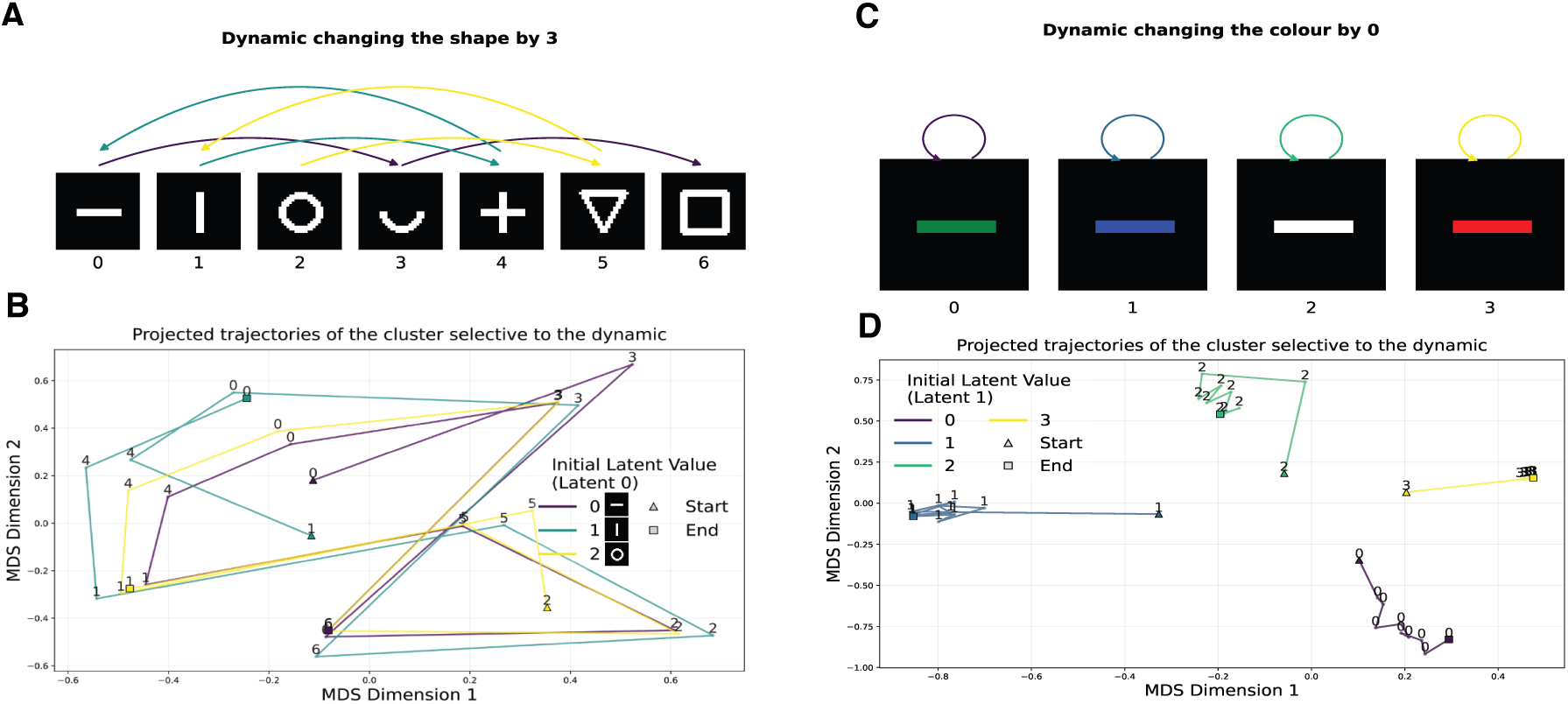
Trajectory of the activity of two clusters. **A** Illustration of the dynamic that changes the shape by 3. **B** Trajectory of the activity of the cluster that is most selective to the dynamic presented above, projected using MDS, using 3 different values at t=0 for the shape latent. For example, when the first shape is the horizontal line, with index 1, adding 3 will result in index 4, which is the plus sign. Adding 3 again results in index 0 the horizontal line, since we perform the addition modulo 7.**C-D** Same as **A-B** but for the dynamic of the colour where it stays equal.

### S7 Testing on the 3D Shapes dataset

To test the how general our results are, we simulated another task typically used in disentanglement literature, the 3D shapes dataset [3, 18] Figure S7. This dataset originally has 6 different latents: the floor hue, wall hue, object hue, scale, shape, and orientation. The hues have 10 possible values, the scale has 8, the shape has 4, and the orientation has 15. We chose to use four latents: the wall and object hue, the shape and the scale, and downsampled their possible values by a factor of 2, as having too much latents and values made it hard for the network to learn the predictive task. We generated sequences the same way we did for our dataset, by starting from randomly sampled latents and adding dynamics to them. We used 3 dynamics per latent, resulting in 81 contexts, from which we used 64 to train the network, and 17 to test compositional generalisation. We used the exact same architecture as the one used for our dataset in the main results. We obtain similar results to the one we had for our dataset. The predictive model perfectly disentangles every latent, (Figure S7 B), and has modular representations of the dynamics (Figure S7 C,D), which allow for compositional generalisation (Figure S7 E). The results for this dataset are more sensitive to the hyperparameter choice, presumably because the task is harder. For the predictive model, we used *λ_weights_* = 0.01*, λ_act_* = 0.1. For the memory and autoencoder, we chose the parameters that maximised the disentanglement score, for the memory model,*λ_weights_* = 0.001*, λ_act_*= 0.001, and for the autoencoder *λ_weights_*= 0.1*, λ_act_*= 0.001.

**Figure S7:**
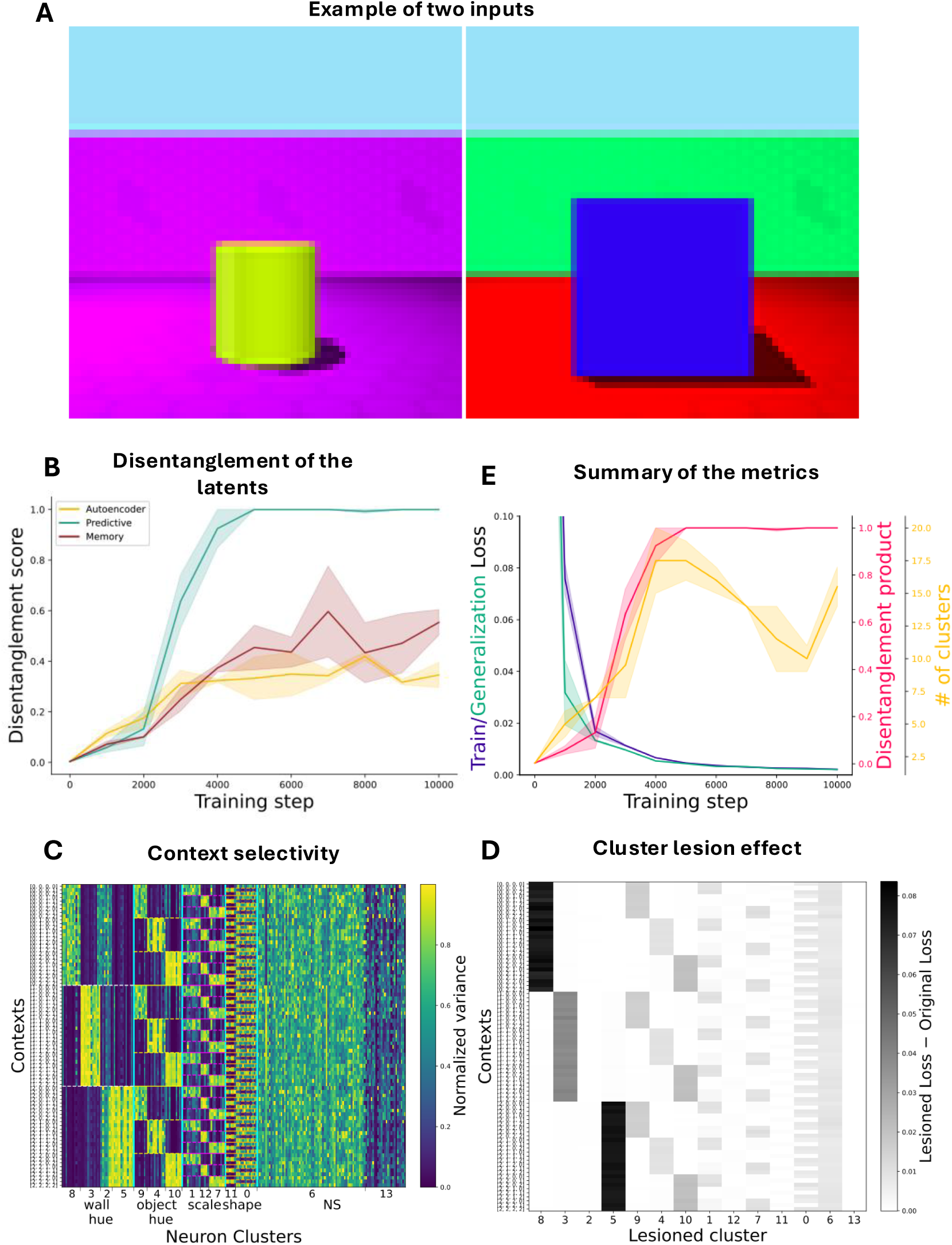
Analysis for the 3D shapes dataset. **A** Examples of two images from the dataset. **B** Disentanglement scores of the three models over training. The predictive model is the only one that disentangles perfectly the 4 latents. **C** Context selectivity profile of the predictive model. Each dynamic has a cluster, except for the third one of the shape. **D** Effect of lesioning a cluster on the reconstruction loss. It is isolated to the sequences for which each cluster is selective to. **E** Recapitulation of the metrics, and showing compositional generalisation by the training and generalisation loss reaching the same point.

## Notes

### Competing Interest Statement

The authors have declared no competing interest.

### Summary of Updates

Added an explanation for the disentanglement metric, and additional simulations to explain the mechanism.

